# DNAscan2: a versatile, scalable, and user-friendly analysis pipeline for next-generation sequencing data

**DOI:** 10.1101/2022.05.12.491669

**Authors:** Heather Marriott, Renata Kabiljo, Ahmad Al Khleifat, Richard J Dobson, Ammar Al-Chalabi, Alfredo Iacoangeli

## Abstract

The current widespread adoption of next-generation sequencing (NGS) in all branches of basic and clinical genetics fields means that users with highly variable informatics skills, computing facilities and application purposes need to process, analyse, and interpret NGS data. In this landscape, versatility, scalability, and user-friendliness are key characteristics for an NGS analysis tool. We developed DNAscan2, a highly flexible, end-to-end pipeline for the analysis of NGS data, which (i) can be used for the detection of multiple variant types, including SNVs, small indels, transposable elements, short tandem repeats and other large structural variants; (ii) covers all steps of the analysis, from quality control of raw data to the generation of html reports for the interpretation and prioritisation of results; (iii) is highly adaptable and scalable as it can be deployed and run via either a graphic user interface for non-bioinformaticians, a command line tool for personal computer usage, or as a Snakemake workflow that facilitates parallel multi-sample execution for high-performance computing environments; (iv) is computationally efficient by minimising RAM and CPU time requirements.

**Availability and Implementation:** DNAscan2 is implemented in Python3 and is available to download as a command-line tool and graphical-user interface at https://github.com/KHP-Informatics/DNAscanv2 or a Snakemake workflow at https://github.com/KHP-Informatics/DNAscanv2_snakemake.

## Introduction

Thanks to its growing accessibility and affordability, next-generation sequencing (NGS) is now being adopted in all fields of clinical and biomedical genetics. As a consequence, a broad audience of users requires flexible and easy-to-use bioinformatics software able to adapt to their informatics proficiency, computing infrastructure and study objectives. Current publicly available NGS pipelines normally focus on the analysis of specific types of genetic variants, e.g. only SNVs and small indels, or only structural variants, do not cover the whole analysis process (i.e. they are not end-to-end), and are not suitable for users with limited informatics skill (Chiang et al., 2015; Collins et al., 2020; DePristo et al., 2011; Zarate et al., 2020). Although, tools that focus on solving some of these factors exist, e.g. being end-to-end (Causey et al., 2018) or user-friendly (Blankenberg et al., 2010), to our knowledge, only commercial bioinformatics solutions which are not accessible to the majority of NGS users, cover all of these aspects (Miller et al., 2015). On such a basis, we developed DNAscan2, a versatile and scalable end-to-end pipeline for the analysis of NGS data. Its graphic user interface (GUI) is suitable for users with any informatics proficiency, whilst its Snakemake implementation makes its analyses highly reproducible, scalable, and adaptable for use on high-performance computing (HPC) facilities. DNAscan2 covers all steps of the analysis process, covering quality controls on the raw data, read mapping, variant calling and annotation of a wide range of genetic variants (SNVs, small indels, and structural variants), results prioritisation and the generation of interactive reports. Finally, DNAscan2 is optimised to provide a comprehensive and reliable data analysis (e.g. by using multiple variant callers) while minimising the computational requirements.

## Results and Implementation

DNAscan2 is written in Python3 and is an open-source software tool available to download from GitHub (https://github.com/KHP-Informatics/DNAscanv2). The installation of its software and database dependencies (Supplementary Table 1) can be performed manually, with a bash helper script or via a GUI. An Anaconda (Anaconda Software Distribution. (2022)) environment file of available binary dependencies is also provided for those who want to install software without package conflicts. The full list of dependencies with their installation specifics are shown in Supplementary Table 2.

### New and Upgraded Features

DNAscan2 maintains the backbone architecture of DNAscan (Fig. 1, Panel A; (Iacoangeli et al., 2019a). Users can supply human DNA NGS data (targeted sequencing, whole exome sequencing (WES) or whole genome sequencing (WGS)) in paired or single-end FASTQ, SAM, BAM, or CRAM format. However, unlike in DNAscan, where users could select one of three modes (fast, normal, and intensive) to tailor the computational requirements of the alignment and variant calling process to their availability, DNAscan2 implements a single protocol that combines a set of aligners and variant callers tailored on the type of variants the user in interested in (see Fig 1 and Supplementary Table 2 for details). For example, DNAscan2 uses HISAT2 (Kim et al., 2019) to align genomic reads if the user only wants to perform SNV calling. An additional BWA-mem realignment (Li, 2013) of soft-clipped and unaligned reads to improve small indel, structural variant (SV) and transposable element detection is also available (Supplementary Table 3). Moreover, the set of variant callers included in DNAscan2 contains multiple SV callers that allow for improved detection of a wider range of SV types. Descriptions of the benchmarking procedure and detailed results for selected SNV/indel and structural variant tools in the following sections are available in the Supplementary Information.

**Fig. 1.**
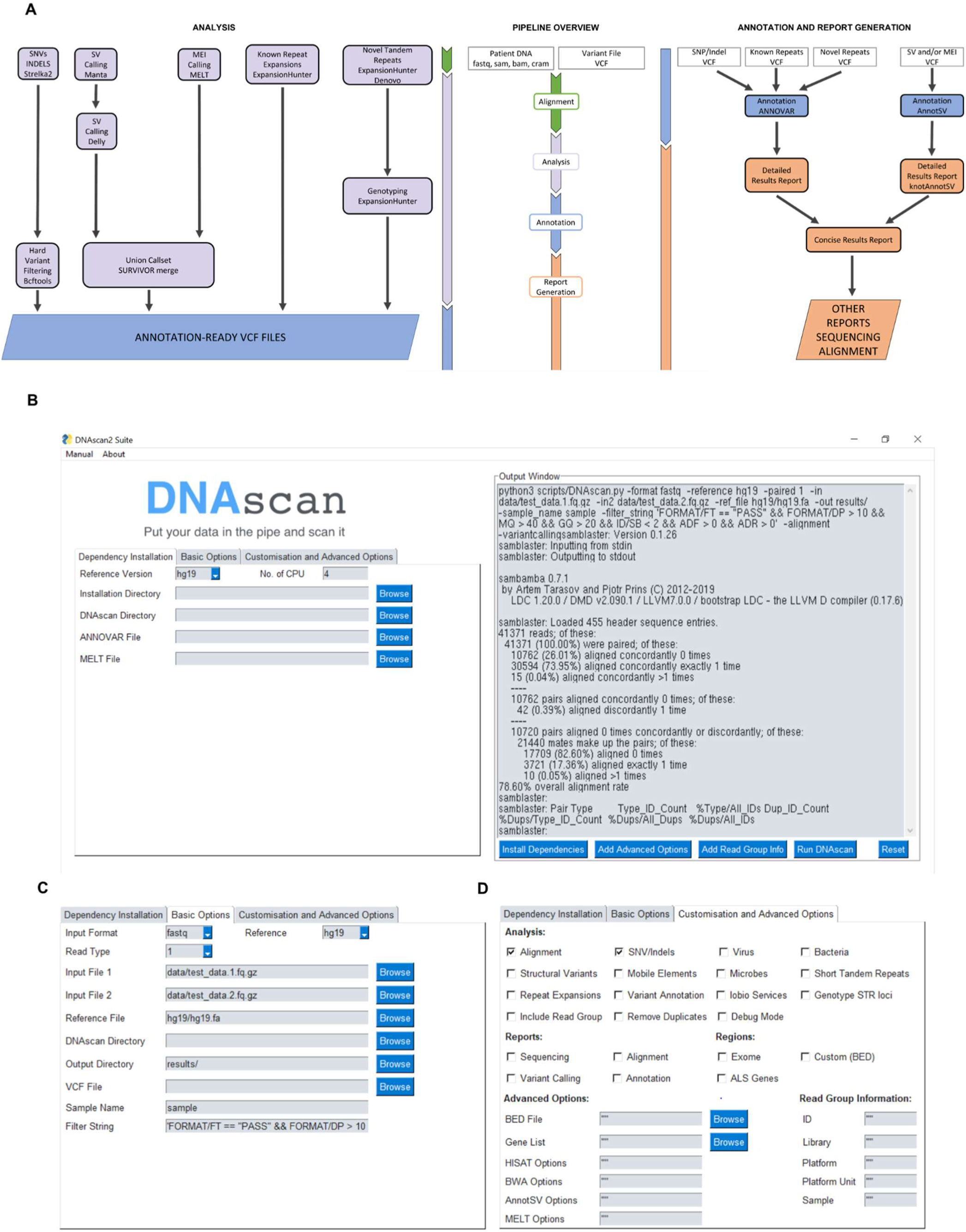
Panel A: Overview of the DNAscan2 pipeline. There have been two major updates to the architecture of the workflow from the original DNAscan version, concerning variant calling and filtering (left) and the annotation and reporting of disease relevant genetic variants (right). Panel B: Screengrab of the main DNAscan2 GUI with an example alignment output, developed using PySimpleGUI and initiated with a Python script supplied with the DNAscan2 source code. Panel C: ‘Basic Options’ tab displaying the default settings of DNAscan2. Panel D: ‘Customisation and Advanced Options’ tab which allows the user to tailor DNAscan2 execution to their requirements.

### SNV and Indel Calling

The Strelka2 small variant caller (Kim et al., 2018) has replaced Freebayes (Garrison and Marth, 2012) and GATK Haplotype Caller (Poplin et al., 2018) for both SNV and indel calling (Fig. 1, Panel A), as it has a similar performance for SNVs and consistently demonstrates a higher precision and F-measure for indel detection on both NA12878 WES and HG002 WGS samples for both standard calls (Supplementary Fig.1, Supplementary Fig. 2A) and medically relevant genetic variants in challenging regions (Supplementary Fig.2B).

### Structural Variant Calling

An enhanced structural variant calling protocol was developed via the addition of Delly (Rausch et al., 2012) to call inversion and deletion variants as well as tandem duplications and translocation events. Delly exhibits a 28-35% higher F-measure for small (50-1000bp) and medium (1001-10000bp) deletions (Supplementary Fig. 3A) on BWA-mem and HISAT2 aligned HG002 WGS reads generated with DNAscan, in addition to a 35% increase in precision for small (101-1000bp) haplotype-resolved inversion calls (Sudmant et al., 2015) on simulated NA12878 WGS reads (Supplementary Fig. 4A). Furthermore, almost all true positive deletion and inversion calls for both datasets are exclusive to Delly or shared by both Manta and Delly (Supplementary Fig. 3B, Supplementary Fig. 4B). This improved calling comes at the expense of increased runtime, with DNAscan2 taking ∼30h longer to run (Supplementary Table 4).

### Transposable Element and Short Tandem Repeat Discovery

The protocol for the detection of mobile element insertions (Alu, SVA and LINE1) and tandem repeats has been substantially improved with the addition of new state-of-the-art tools. Mobile elements can now be discovered and genotyped via MELT (Gardner et al., 2017) and a genome-wide non-reference short tandem repeat loci profile with details of the motif composition and estimated repeat size of each identified repeat can be generated using ExpansionHunter Denovo (Dolzhenko et al., 2020). Users also have the option to convert the repeat loci into a catalogue format compatible with ExpansionHunter (using a conversion script available at https://github.com/francesca-lucas/ehdn-to-eh) to undergo repeat size estimation (Fig. 1, left panel).

### Variant Annotation and Report Generation

The spectrum of variant calls that can now be annotated has been extended to include structural and transposable elements (Fig.1., right panel) with the incorporation of AnnotSV (Geoffroy et al., 2018), in addition to known and novel repeat expansions using user-defined ANNOVAR databases. Additionally, an HTML report of variants annotated with AnnotSV produced with the knotAnnotSV program (Geoffroy et al., 2021), and a generalised annotation report giving the type, genomic location, overlapping genes and population variant frequency of all identified variants are created for the user’s convenience (Supplementary Fig. 5).

### Snakemake and GUI Accessibility

To expand the accessibility of DNAscan2, both a graphical user interface (Fig 1. Panels B, C, D) and a Snakemake workflow (available at https://github.com/KHP-Informatics/DNAscanv2_snakemake) have been developed. This renders DNAscan2 available as both an easy-to-use, end-to-end program via its GUI and as a scalable command line tool which can be executed on high-performance computing facilities.

### Computational Performance

DNAscan2 is optimised to minimise the computational resources necessary for its use. The average memory usage in the SNV and indel calling stage for WGS is approximately 1Gb (Supplementary Table 4, Supplementary Fig. 6); an improvement of 97% compared with DNAscan. DNAscan2 can complete the full protocol, including full SV calling and annotation, on WGS data in 50 hours using 4 CPUs and ∼15Gb RAM (Supplementary Table 4, Supplementary Fig. 6, Supplementary Fig. 7), much within the hardware specifications of a midrange personal computer.

## Conclusions

DNAscan2 adapts to the heterogenic needs of the wide audience using NGS data nowadays. It shows potential to be of great value for a broad range of users and uses, e.g. clinical geneticists focusing on disease diagnostics (Iacoangeli et al., 2019b; Supplementary Fig. 8, Supplementary Table 5), as well as biomedical researchers working on large-scale genomic studies.

## Acknowledgements

The authors acknowledge use of the research computing facility at King’s College London, Rosalind (https://rosalind.kcl.ac.uk), which is delivered in partnership with the National Institute for Health Research (NIHR) Biomedical Research Centres at South London & Maudsley and Guy’s & St. Thomas’ NHS Foundation Trusts, and part-funded by capital equipment grants from the Maudsley Charity (award 980) and Guy’s & St. Thomas’ Charity (TR130505). The views expressed are those of the author(s) and not necessarily those of the NHS, the NIHR, King’s College London, or the Department of Health and Social Care.

