## Supplementary Information for "DNAscan2: a versatile, scalable, and user-friendly analysis pipeline for next-generation sequencing data"

12/05/2022

**Supplementary Materials and Methods**

**Datasets**

To compare the SNV and indel calling performance of Freebayes, Strelka2 and GATK Haplotype Caller, we used the paired-end Illumina HiSeq4000 WES of NA12878 (NCBI SRA Accession: ERR1905890) and Illumina HiSeqX PCR-free WGS of HG002 (NCBI SRA Accession: SRR14724544). Samples were aligned to the hg19 genome using the -alignment flag of DNAscan in both fast and normal mode prior to SNV and indel calling. Alignment is performed by HISAT2 in fast and normal mode, with BWA-mem introduced in the latter for realignment of soft clipped and unaligned reads to improve the detection of small indels and structural variants in downstream calling steps. Structural variant calling performance of Manta and Delly was evaluated using the aligned HG002 reads in addition to simulated paired-end Illumina WGS reads containing the positions of haplotype-resolved deletion and inversion calls of NA12878 (Sudmant *et al.*, 2015), generated and aligned to the hg38 genome using the VISOR package (Bolognini *et al.*, 2020).

For comparing the variant calling and computational power of the new and previous implementations of DNAscan, we utilised 10 UK control WGS samples sequenced as part of the Project MinE amyotrophic lateral sclerosis sequencing consortium (Project MinE ALS Sequencing Consortium, 2018). Genomic DNA from venous blood drawn from patients and controls was isolated using standard methods before DNA integrity was assessed using gel electrophoresis. Samples were sequenced using Illumina’s FastTrack Services (San Diego, CA) on the Illumina HiSeq2000 platform. PCR-free library preparation was used to perform 100bp paired-end sequencing, which yielded ~40x coverage across each sample. Sequenced samples were aligned to the hg38 genome using BWA-mem and were provided to us in CRAM format.

**Benchmarking of variant callers to include in DNAscan2**

It was not necessary for callers associated with every additional functionality to undergo benchmarking as they either demonstrate beneficial value for identifying novel disease-relevant loci i.e. ExpansionHunter Denovo (Rafehi *et al.*, 2019; Fazal *et al.*, 2020), or consistently show high performance in benchmarking studies i.e. MELT for MEI detection (Kosugi *et al.*, 2019; Vendrell-Mir *et al.*, 2019).

**SNVs and Indels**

The calling performance of Freebayes and GATK HaplotypeCaller was assessed by running DNAscan with the -variantcalling flag in fast, normal and intensive modes for the NA12878 WES and HG002 WGS samples. Fast and normal mode uses Freebayes to call SNVs and indels from a coordinate sorted BAM file. Intensive mode employs GATK HaplotypeCaller to call indels from genomic positions (identified from HISAT2 and BWA-mem alignment) that contain at least one read that potentially harbours a deletion or insertion variant. A similar approach was adopted for Strelka2; both SNVs and indels were called from aligned reads generated by fast and normal mode of DNAscan, with the BED file of probable indel variant positions being used to separate output into SNV and indel variant files with the VCFtools --exclude-bed and --bed flags, analogous to DNAscan intensive mode. For NA12878, Strelka2 was run with the --exome flag.

Benchmarking of variant calling performance was performed using hap.py version 0.3.14 (*Haplotype Comparison Tools*, 2021) with the RTG vcfeval engine. An example command is shown below, where the baseline VCF and confident regions BED file correspond to the appropriate true positive callsets used to evaluate the SNV and indel calls:

hap.py --false-positives {confident_regions.bed} –-reference hg19.fa –-report-prefix {outdir} –-engine=vcfeval –-threads {4} {baseline.vcf} {comparison.vcf}

The NA12878 WES calls were evaluated against the National Institute of Science and Technology (NIST) with Platinum Genomes phase transfer small variant truth set (version 3.3.2). Confident call regions were defined as the hg19 exome probe locations from the Agilent SureSelect Human All Exon v5 kit (ELID:S04380110). HG002 calls were evaluated with both the truth variants and confident regions of the NIST small variant benchmarking (version 4.2.1) and Genome In A Bottle Challenging Medically-Relevant Genes (GIAB CMRG version 1.00) datasets.

**Structural Variants**

The structural variant detection power of Manta and Delly was assessed in a similar fashion to that of SNV and indel callers. The HISAT2 and/or BWA-mem aligned BAM files of HG002 and NA12878 WGS samples were processed by Manta as per the old DNAscan implementation, with the same approach adopted for Delly as to obtain a comparable dataset.

Calls were then evaluated using Truvari version 2.2.1 (*spiralgenetics/truvari*, 2021) using the following command:

truvari bench -b {baseline.vcf} -c {comparison.vcf} -f {reference.fasta} -o {outdir} --includebed {confident_regions.bed}

For HG002, the performance of Delly and Manta on detecting deletions was evaluated using the variant calls and confident regions of the GIAB CMRG deletion SV truth set (version 0.6). Deletion and inversion calls of NA12878 were evaluated using calls from the 1000 Genomes structural variant map (Sudmant *et al.*, 2015) as the comparison VCF. To further compare the obtained structural variant calls, the --sizemin and --sizemax flags were supplied to Truvari. Parameters were defined as 50-100, 101-1000, 1001-10000 and 10000-50000bp (the default --sizemax parameter of Truvari).

**Comparison between DNAscan and DNAscan2**

Once DNAscan2 had been finalised and implemented, both versions of DNAscan were run with the -variantcalling, -SV, -expansion, -annotation and -resultsreport flags on each of the 10 Project MinE UK control samples. DNAscan2 was additionally run with the -MEI and -STR flags.

**Performance Metrics**

**Variant Calling Performance**

Both SNV/indel and structural variant calls were evaluated using precision, recall and F-measure metrics calculated by hap.py and Truvari, where precision is $\frac{True Positives}{(True Positives+False Negatives)}$, recall is $\frac{True Positives}{(True Positives+False Positives)}$ and F1 score is $\frac{True Positives}{True Positives+\frac{1}{2}(False Positives+False Negatives)}$ .

Where possible, calling performance of DNAscan and DNAscan2 was assessed by averaging the number of SNV/indel, SV, repeat expansion and MEI calls and STR loci obtained for all Project MinE samples. SV and MEI calls were also separated into their respective subclasses i.e. deletions, insertions, inversions, duplications, Alu, SVA, LINE1. Additionally, the number and percentage of filtered synonymous and non-synonymous SNV variants was acquired from the classification given by refGene annotation.

**Computational Performance**

Computational efficiency of the SNV/indel and structural variant callers was assessed by obtaining the wall and CPU time (in hh:mm:ss format) and RAM usage (maximum resident set size in gigabytes) for HG002 and NA12878 samples from the SLURM sacct command --format “Elapsed,CPUTime,MaxRSS”. The same approach was applied for the comparison of DNAscan and DNAscan2, albeit with time and memory usage being averaged across the 10 samples, and estimated disk read and write usage obtained with addition of “MaxDiskRead,MaxDiskWrite” to the above sacct command.

**Storage Requirements of DNAscan2**

Total storage required for installation of dependencies, databases and references with DNAscan2 for both hg19 and hg38 genome versions was calculated with the du -sh shell command.

**Hardware**

All tests were performed on an Intel Xeon E5-2670 2.6GHz processor with 4 CPUs and 16Gb RAM (in line with the standard computational requirements of DNAscan), except for processes which involved Freebayes, which ran with 4 CPUs and 64Gb RAM.

**Availability of Data and Materials**

The variant call file containing haplotype-resolved deletion and inversion calls of NA12878 is available at:

http://ftp.1000genomes.ebi.ac.uk/vol1/ftp/phase3/integrated_sv_map/supporting/GRCh38_positions/ALL.wgs.integrated_sv_map_v1_GRCh38.20130502.svs.genotypes.vcf.gz

The NA12878 NIST with Platinum Genomes phase transfer small variant callset (version 3.3.2) is available at:

https://ftp-trace.ncbi.nlm.nih.gov/giab/ftp/release/NA12878_HG001/latest/GRCh37/HG001_GRCh37_GIAB_highconf_CG-IllFB-IllGATKHC-Ion-10X-SOLID_CHROM1-X_v.3.3.2_highconf_PGandRTGphasetransfer.vcf.gz

Probe regions used for the whole exome sequencing of NA12878 are available from the Agilent SureDesign Dashboard (https://earray.chem.agilent.com/suredesign/). Account registration and sign-in is required to access the regions file from the Agilent catalogue (S04380110_Regions.bed).

The HG002 NIST small variant benchmarking callset containing variant calls and confident regions files (version 4.2.1) are available at the following directory:

https://ftp-trace.ncbi.nlm.nih.gov/giab/ftp/release/AshkenazimTrio/HG002_NA24385_son/NISTv4.2.1/GRCh37/

The HG002 GIAB CMRG small variant calls and confident regions files (version 1.00) are available at the following directory:

https://ftp-trace.ncbi.nlm.nih.gov/ReferenceSamples/giab/release/AshkenazimTrio/HG002_NA24385_son/CMRG_v1.00/GRCh37/SmallVariant/

**Supplementary Figures**

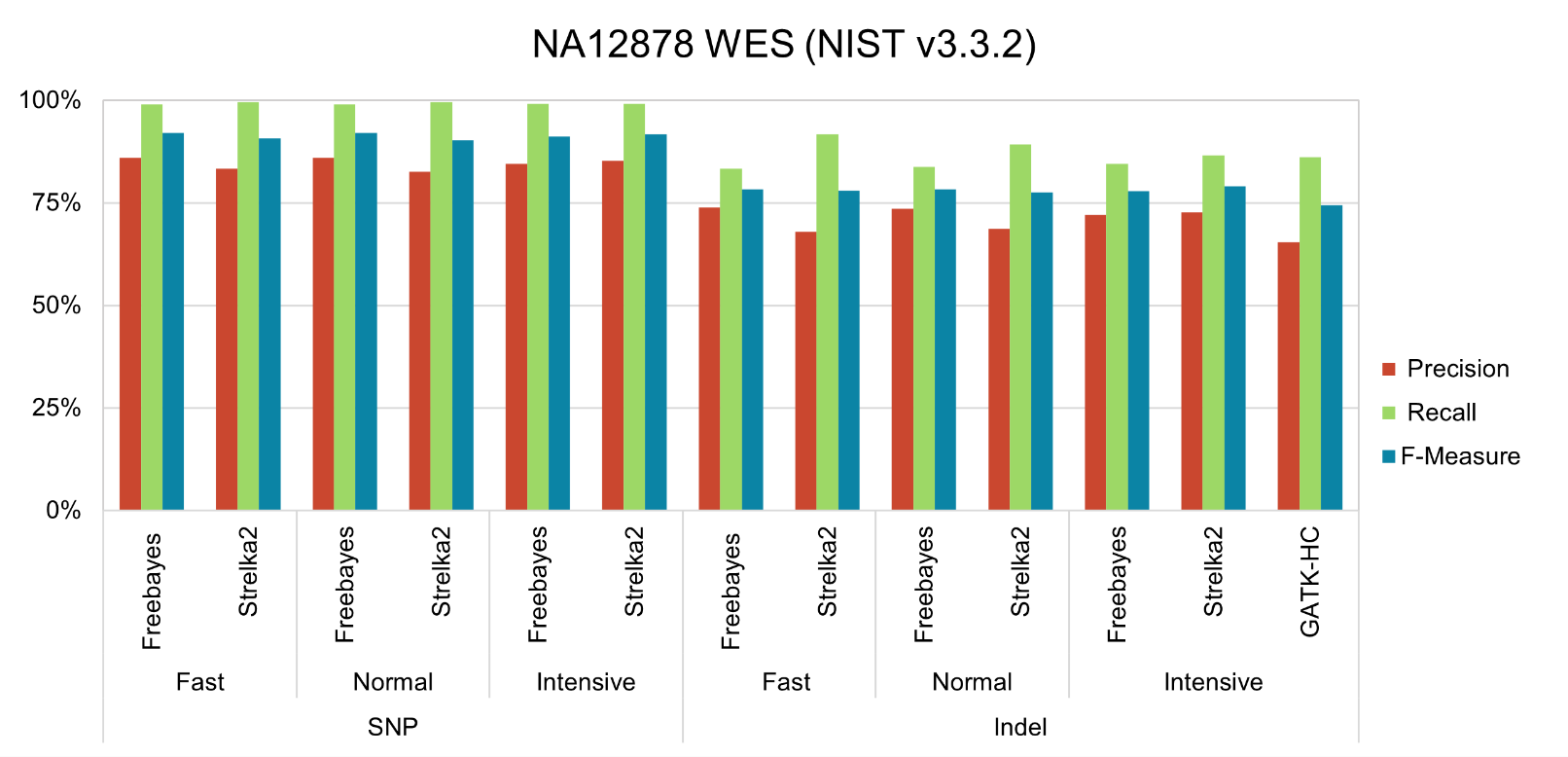

**Supplementary Fig. 1.** SNV and indel calling performance of Freebayes, Strelka2 and GATK HaplotypeCaller on the whole exome sequencing of NA12878, against the NIST small variant benchmarking callset (version 3.3.2). DNAscan was run in fast, normal and intensive mode.

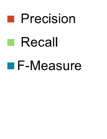

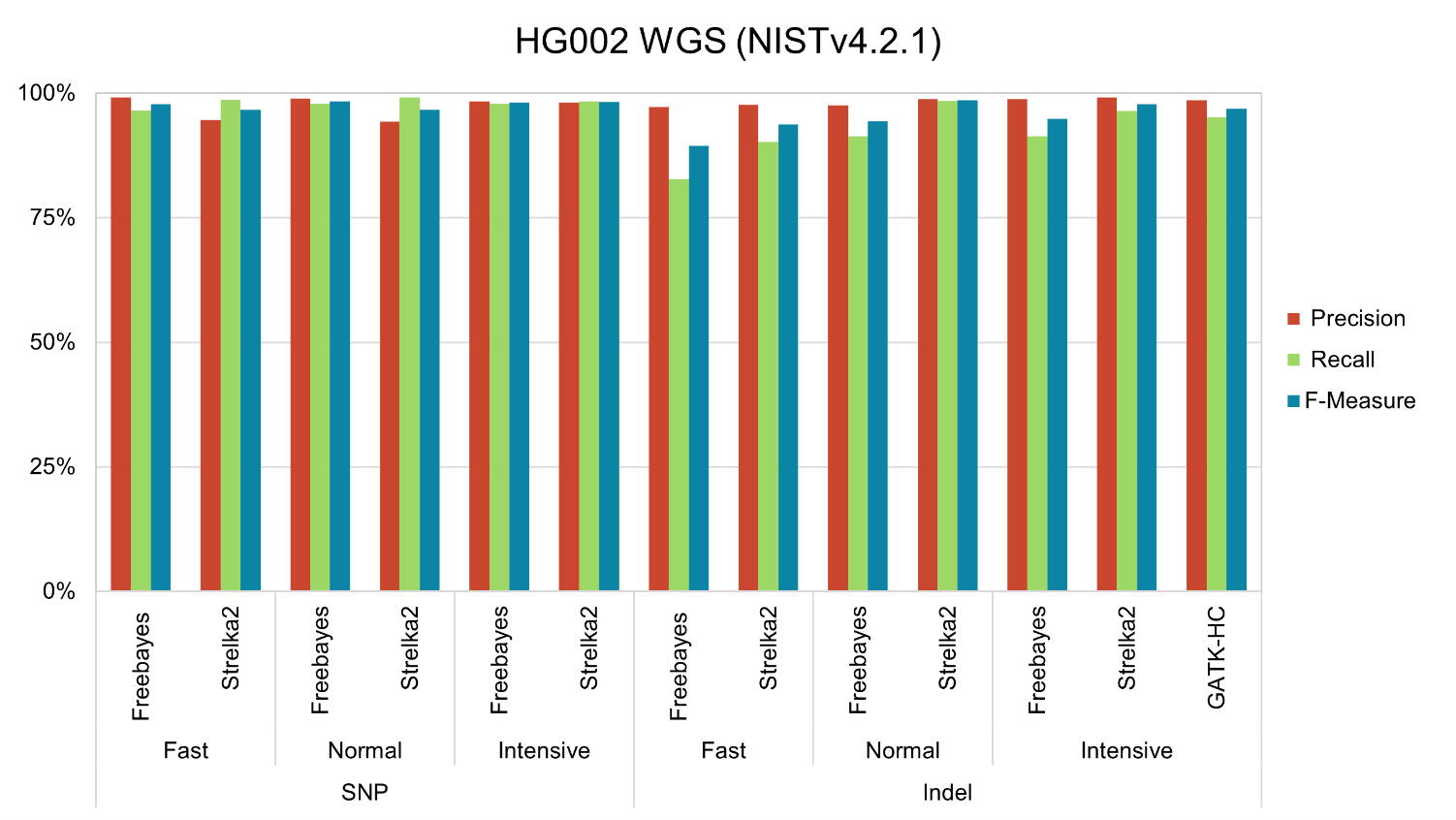

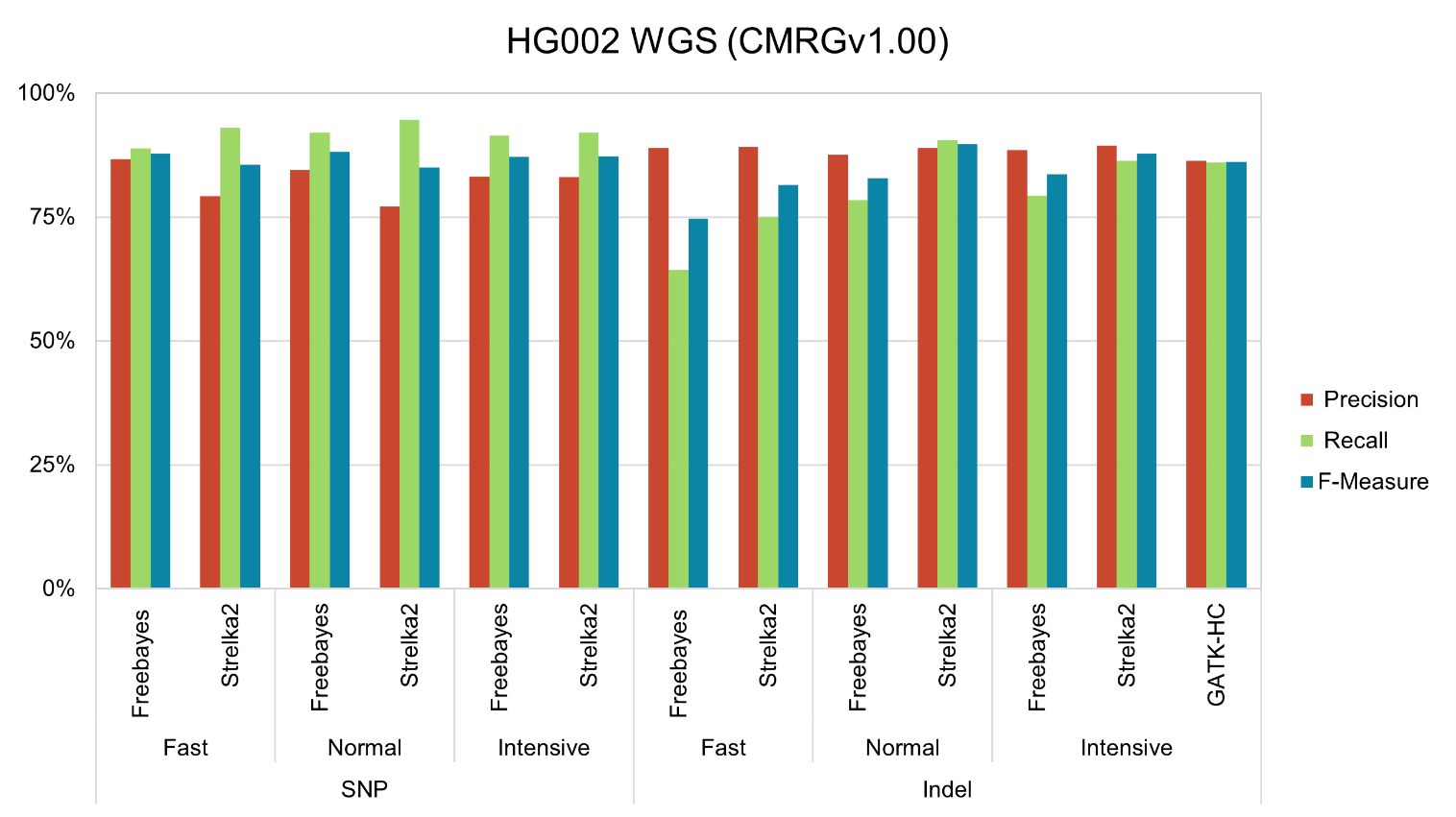

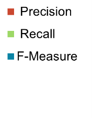

**Supplementary Fig. 2.** SNV and indel calling performance of Freebayes, Strelka2 and GATK HaplotypeCaller on the whole genome sequencing of HG002, against the **A.** NIST small variant benchmarking callset (version 4.2.1), and the **B.** GIAB CMRG structural variant callset (version 1.00). DNAscan was run in fast, normal and intensive mode.

**B**

**A**

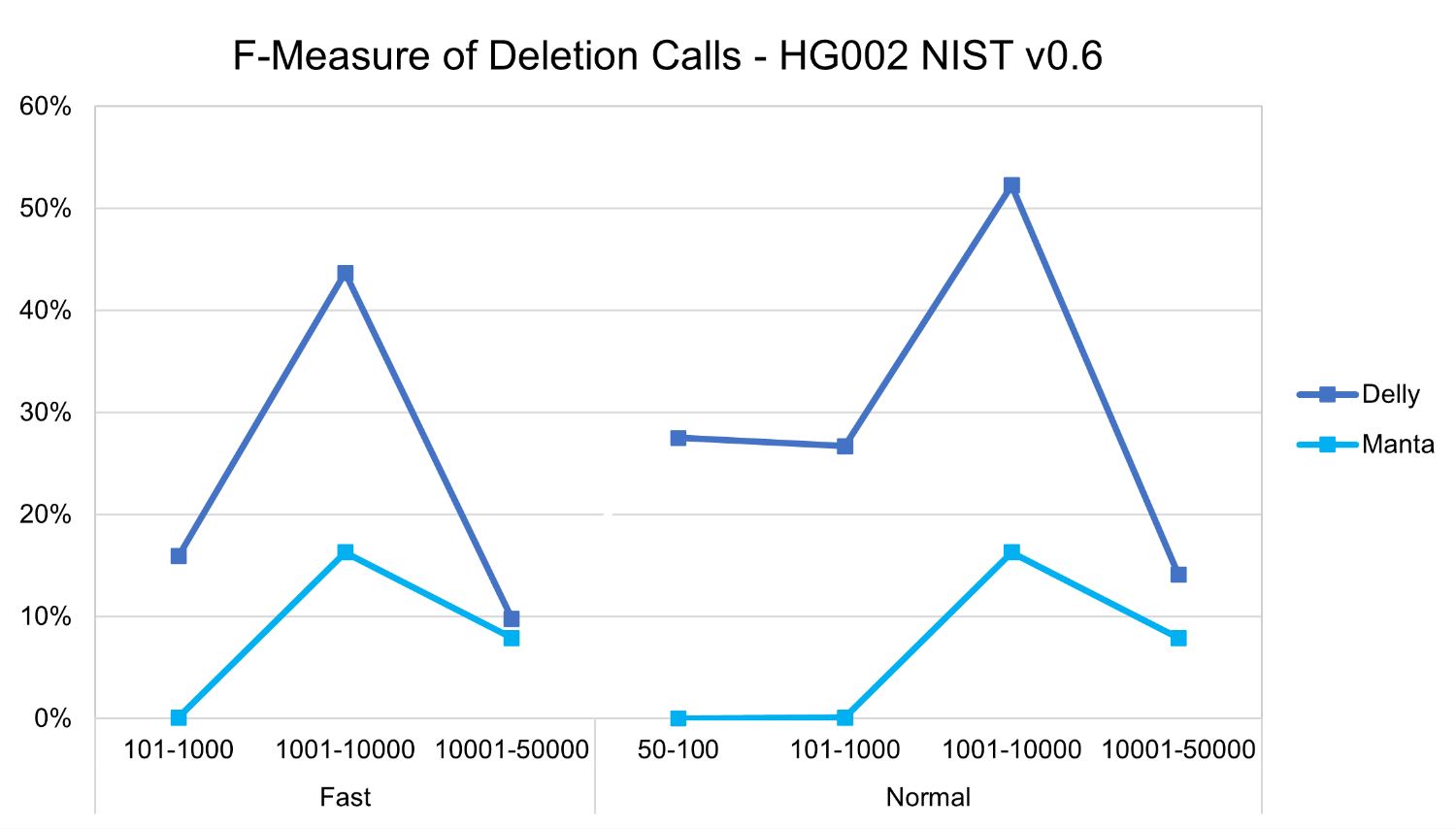

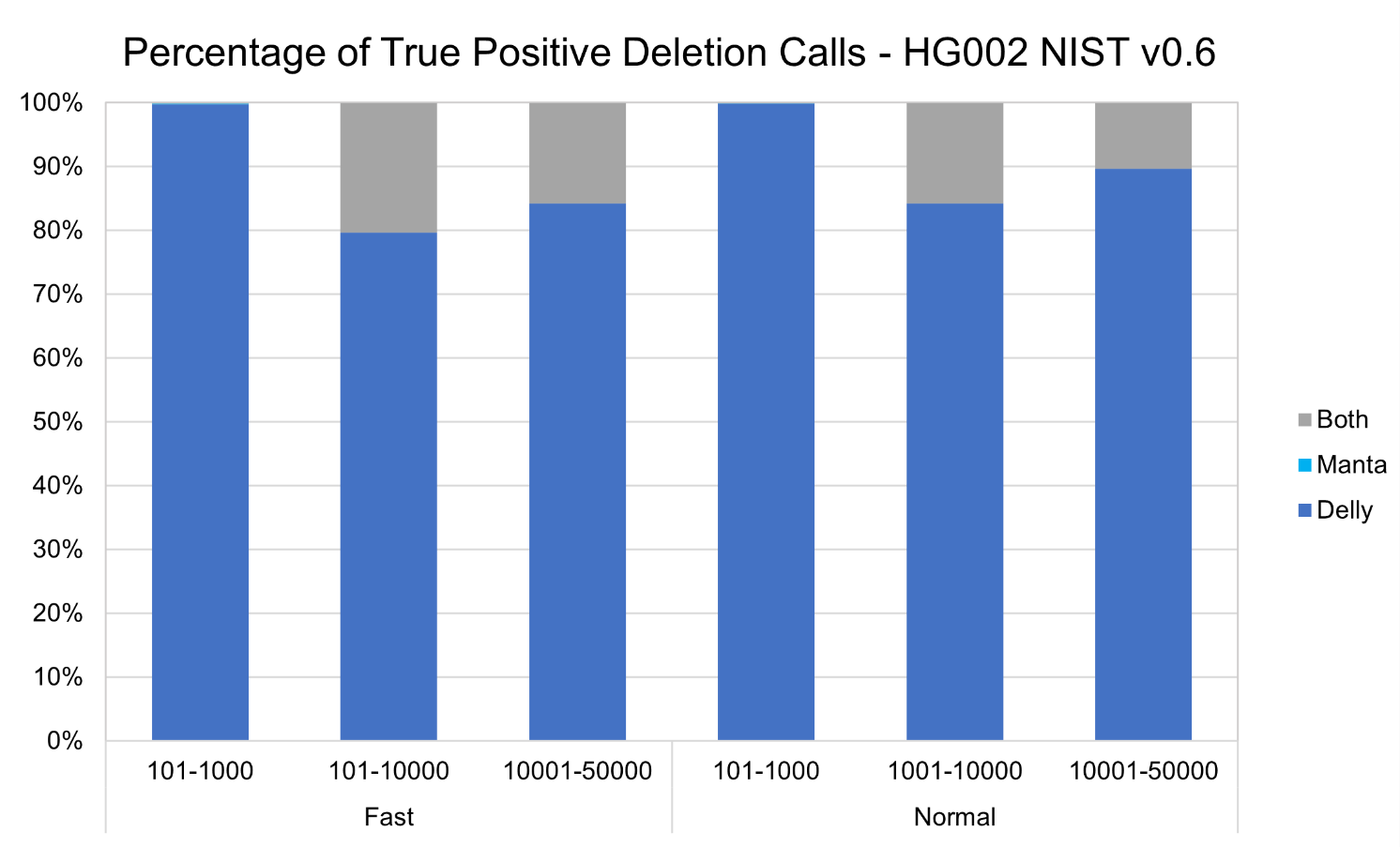

**A**

**B**

**Supplementary Fig. 3.** Deletion SV calling performance of Manta and Delly on the whole genome sequencing of HG002 against the NIST SV benchmarking callset (version 0.6). Fast and normal modes refer to the alignment specifics as defined by DNAscan. Fast: HISAT2 alignment. Normal: HISAT2 alignment and BWA-mem realignment of soft-clipped and/or unaligned reads. **A.** F-Measure of calls, and **B.** Percentage of true positive calls shared by or exclusive to Manta and Delly, in fast and normal mode for multiple deletion variant sizes.

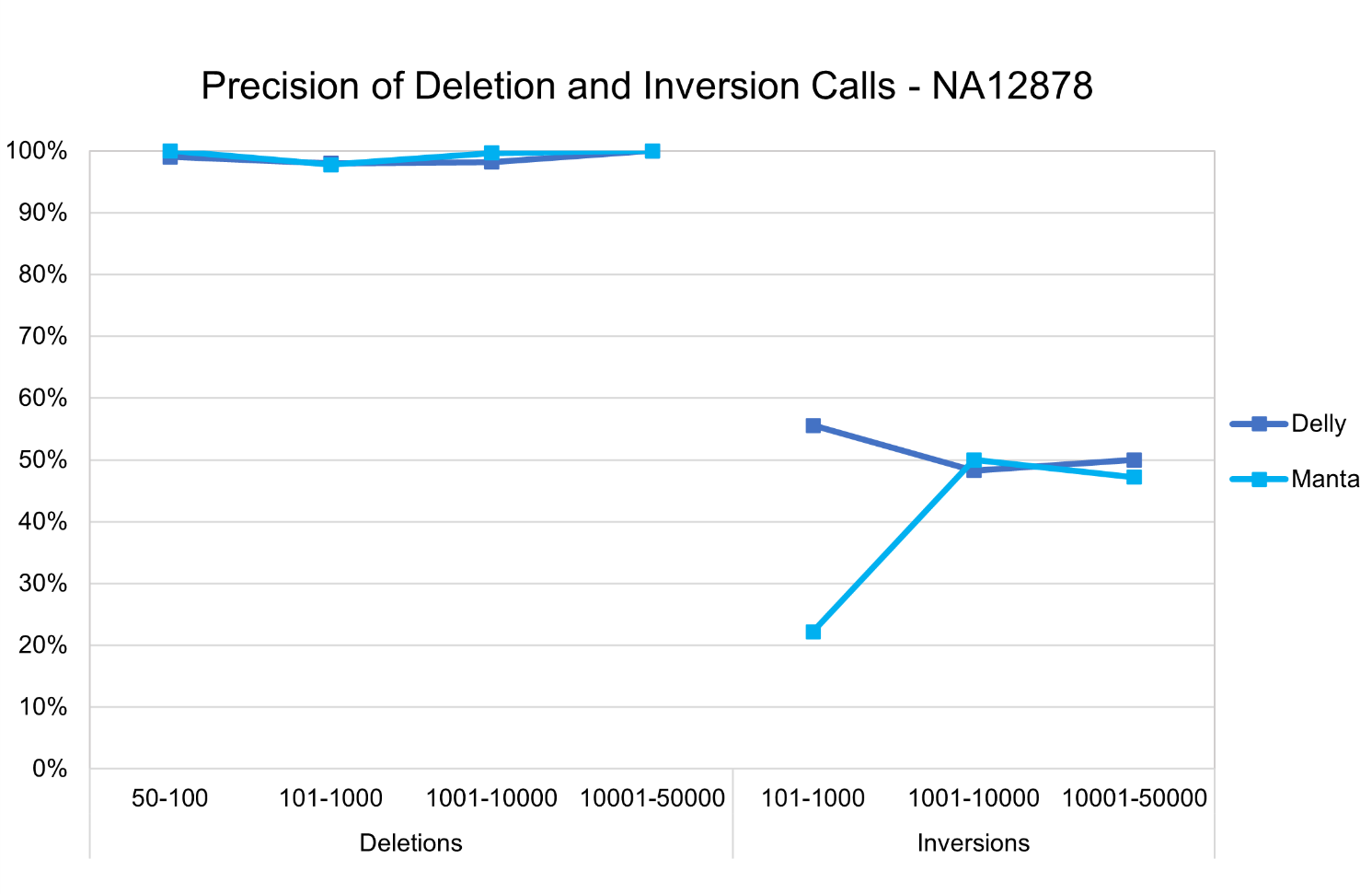

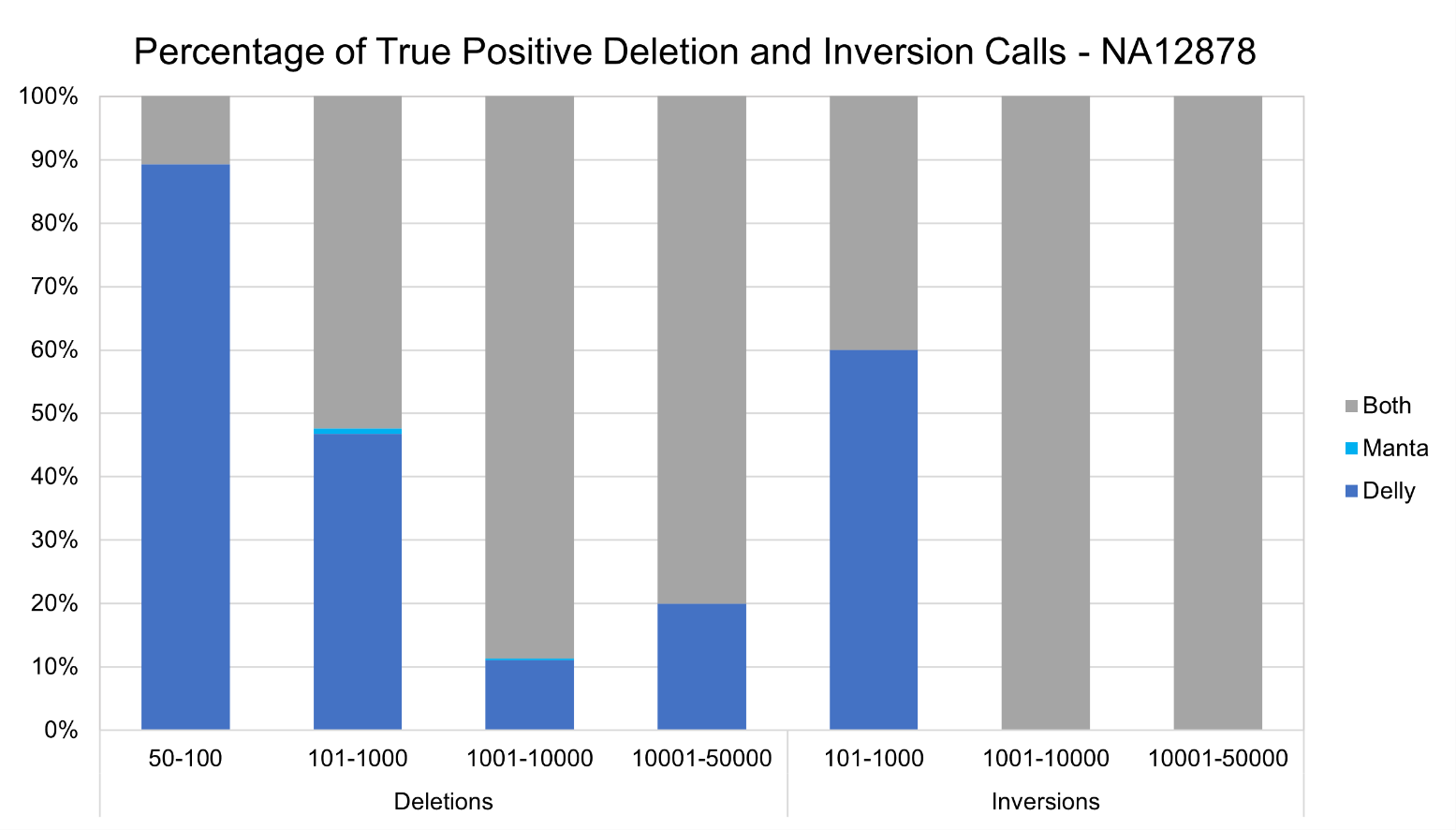

**A**

**B**

**Supplementary Fig. 4.** Deletion and inversion SV calling performance of Manta and Delly on simulated whole genome sequencing reads of NA12878 generated with VISOR. **A.** Precision of calls, and **B.** Percentage of true positive calls shared by or exclusive to Manta and Delly, for multiple deletion and inversion variant sizes.

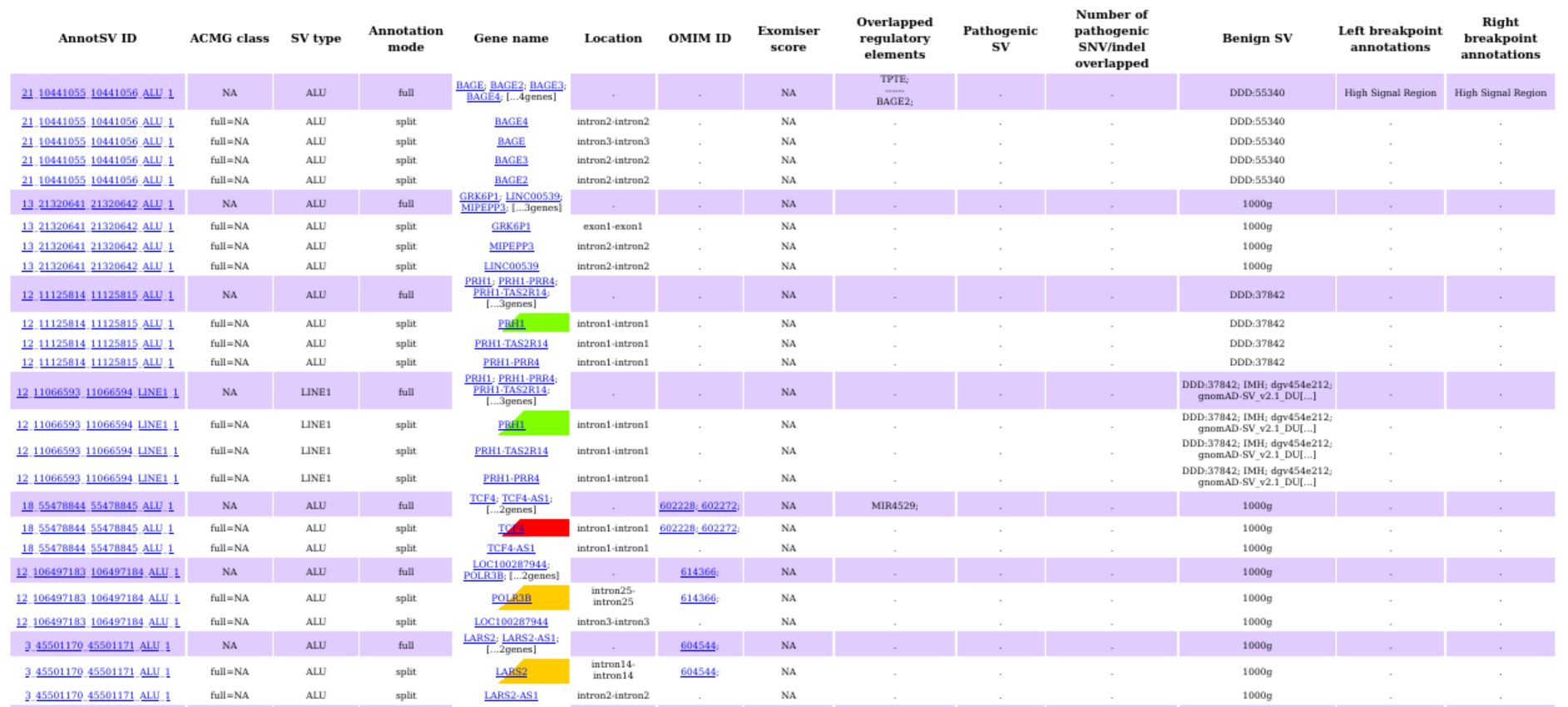

**Supplementary Fig. 5.** Screengrab of the transposable element report generated by DNAscanv2 using knotAnnotSV for 1 Project MinE control sample.

**Supplementary Fig. 6.** Comparison of memory usage (in gigabytes) between DNAscan and DNAscan2 for each step following alignment, categorised by the main stage of the workflow (Analysis, Annotation, Report Generation).

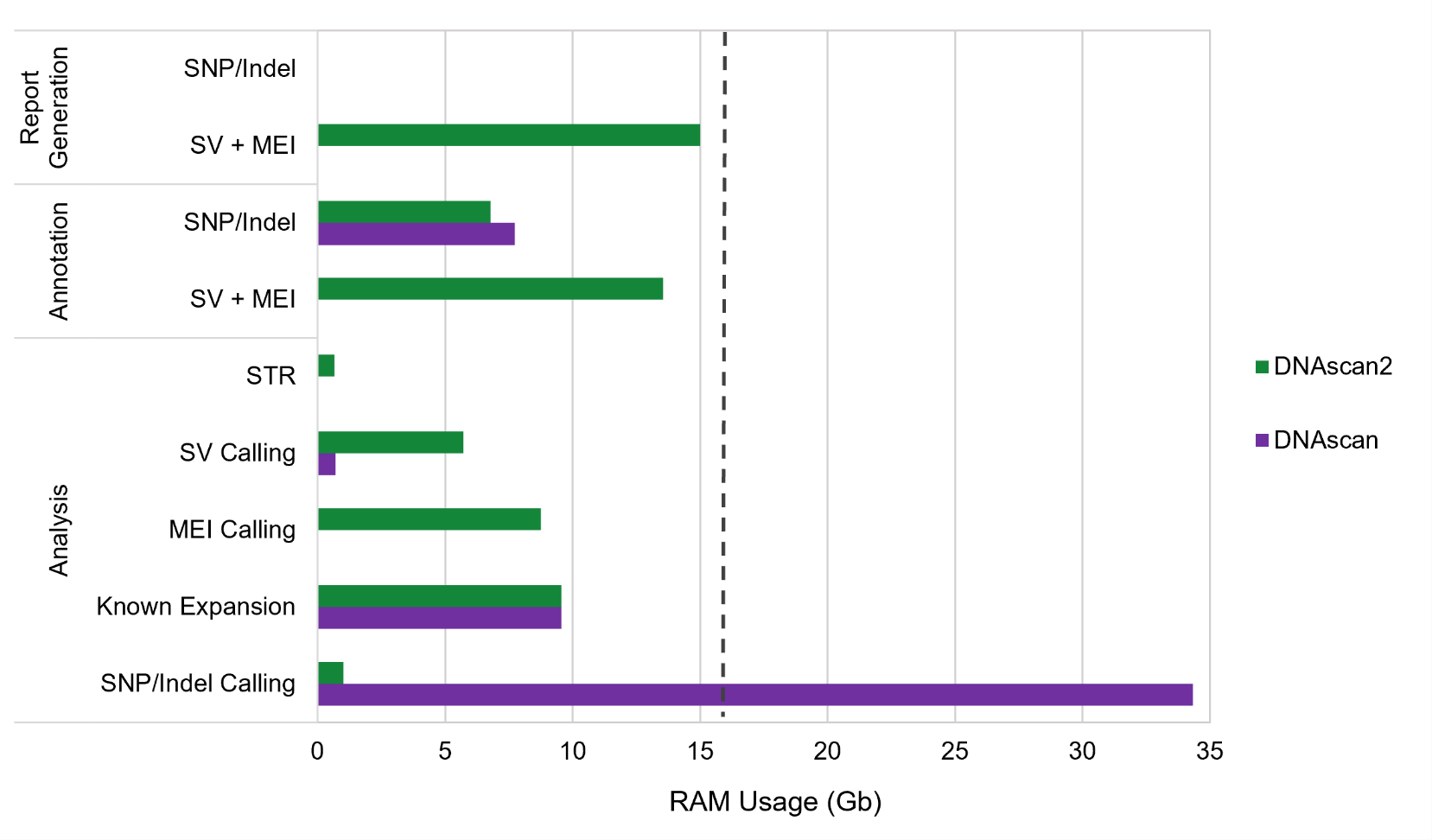

**Supplementary Fig. 7.** Cumulative elapsed time (in hours:minutes:seconds format) between DNAscan and DNAscan2 for each step following alignment, categorised by the main stage of the workflow (Analysis, Annotation, Report Generation). Separate performance metrics for each stage are listed in Supplementary Table 5.

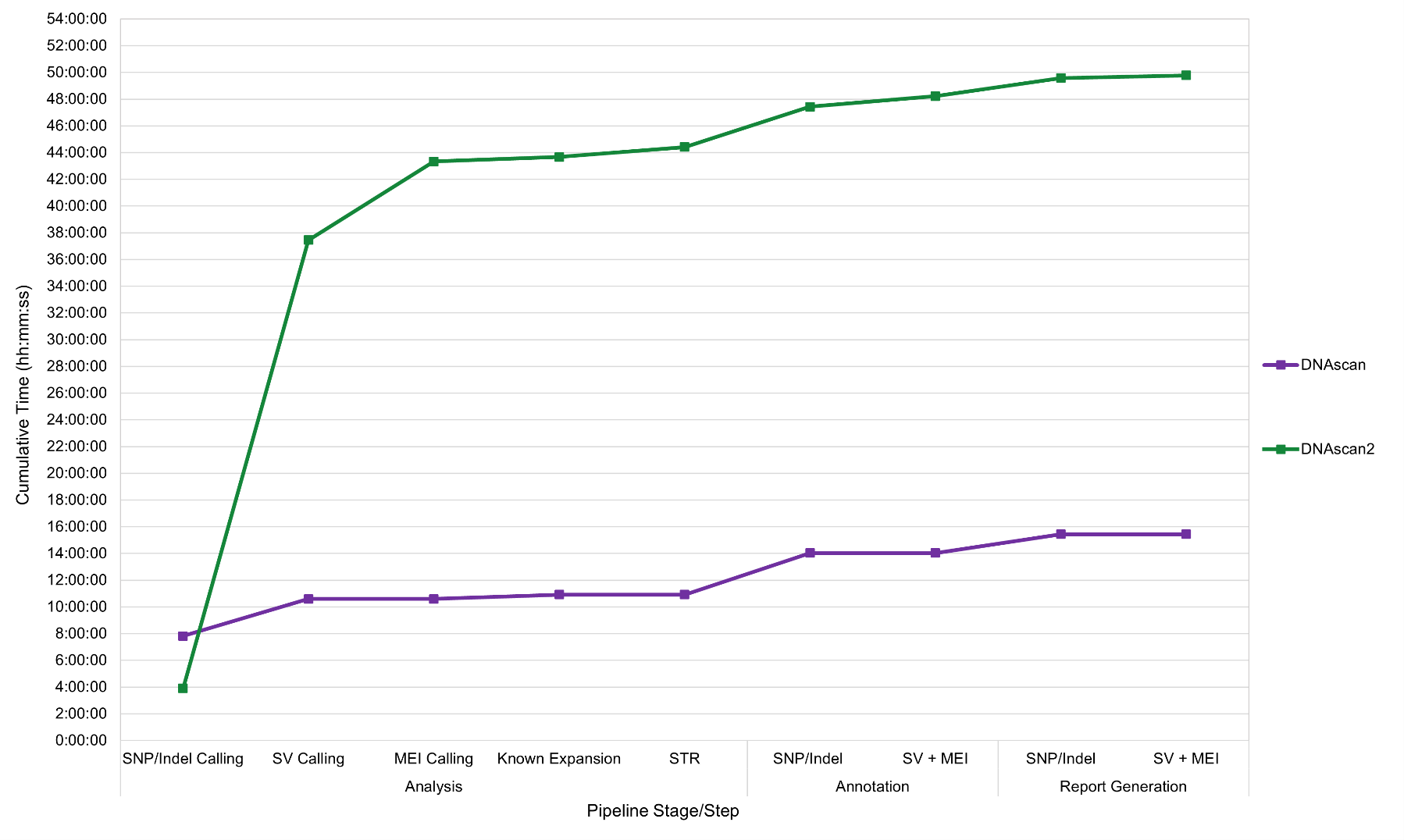

**Supplementary Fig. 8.** The ALSgeneScanner scoring module of DNAscan2. After the main DNAscan pipeline has completed, ALSgeneScanner takes SNV and indel annotation files as input and keeps only the variants whose genes are identified to be associated with ALS (171 in total, from three different sources). Afterwards, each of these filtered variants are scored based on the pathogenicity prediction assigned by each of the 13 prediction databases (SIFT, Polyphen-2 HDIV, PolyPhen-2 HVAR, LRT, MutationTaster, MutationAssessor, FATHMM, PROVEAN, FATHMM-mkl coding, MetaSVM, CADD and Intervar) and then ranked based on the total cumulative score from all programs.

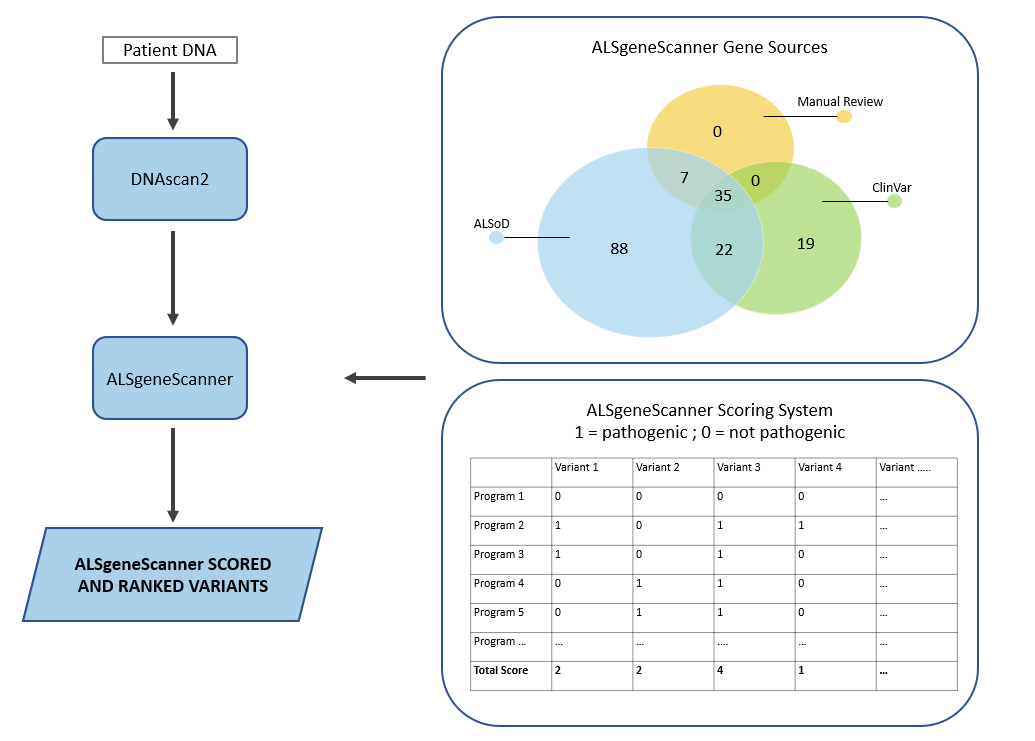

**Supplementary Tables**

| Dependency Category | Required Storage (Gb) | |
| --- | --- | --- |
|  | **hg19** | **hg38** |
| *References* |  |  |
| Reference Genome and Index | 3.16 | 3.27 |
| BWA-mem Index | 5.42 | 5.62 |
| HISAT2 Index | 4.37 | 4.66 |
| *Databases and Catalogues* |  |  |
| ExpansionHunter Variant Catalogue | < 0.01 | < 0.01 |
| ANNOVAR Databases* | 91.34 | 91.67 |
| AnnotSV Databases** | 3.31 | 3.31 |
| *Tools* |  |  |
| Conda Installation | 0.10 | 0.10 |
| Conda-Installed Tools | 1.50 | 1.50 |
| Manually Installed Tools | 12.01 | 12.01 |
| *Additional Disk Space**** | 20.00 | 20.00 |
| Total free space needed: | **141.21** | **142.14** |

**Supplementary Table 1.** Storage requirements of all dependencies necessary to run DNAscan2, divided by reference version. *Default DNAscan2 ANNOVAR databases (refGene, dbnsfp33a, clinvar_20210501, intervar_20180118, avsnp147, exac03,1000g2015aug, gnomad211_genome) were used. **AnnotSV was only run using databases that do not require a commercial licence to download. ***This refers to the additional overhead storage space that MELT requires to temporarily hold intermediate files.

| Pipeline Stage | Tool | Tool Version | Installation Method | Reference | |
| --- | --- | --- | --- | --- | --- |
| *Alignment* |  |  |  | |  |
|  | BWA-mem | 0.7.17 | Conda | | (Li, 2013) |
|  | HISAT2 | 2.2.1 | Conda | | (Kim *et al.*, 2019, p. 2) |
|  | Samblaster | 0.1.26 | Conda | | (Faust and Hall, 2014) |
|  | Sambamba | 0.7.1 | Conda | | (Tarasov *et al.*, 2015) |
| *Analysis* |  |  |  | |  |
|  | Strelka2 | 2.9.10 | Manual Download | | (Kim *et al.*, 2018, p. 2) |
|  | Manta | 1.6.0 | Manual Download | | (Chen *et al.*, 2016) |
|  | Delly | 0.8.3 | Conda | | (Rausch *et al.*, 2012) |
|  | MELT | 2.2.2 | Manual Registration and Download | | (Gardner *et al.*, 2017) |
|  | ExpansionHunter | 3.2.2 | Conda | | (Dolzhenko *et al.*, 2019) |
|  | ExpansionHunter Denovo | 0.9.0 | Manual Download | | (Dolzhenko *et al.*, 2020) |
|  | SURVIVOR | 1.0.7 | Manual Download | | (Jeffares *et al.*, 2017) |
| *Annotation* |  |  |  | |  |
|  | ANNOVAR | 08/06/2019 | Manual Registration and Download | | (Wang, Li and Hakonarson, 2010) |
|  | AnnotSV | 3.0.9 | Manual Download | | (Geoffroy *et al.*, 2018) |
| *Report Generation* |  |  |  | |  |
|  | FastQC | 0.11.9 | Conda | | (*Babraham Bioinformatics - FastQC A Quality Control tool for High Throughput Sequence Data*, no date) |
|  | MultiQC | 1.10.1 | Conda | | (Ewels *et al.*, 2016) |
|  | knotAnnotSV | 1.0.0 | Manual Download | | (Geoffroy *et al.*, 2021) |
| *General* |  |  |  | |  |
|  | Miniconda | 4.10.3 | Manual Download | | (Anaconda Software Distribution. (2022)) |
|  | Perl | 5.26.2 | Conda | |  |
|  | Python | 3.8 | Conda | |  |
|  | Biopython | 1.78 | Conda | | (Cock *et al.*, 2009) |
|  | Pysam | 0.16.0.1 | Conda | | (*Pysam*, 2021) |
|  | SAMtools | 1.9 | Conda | | (Li *et al.*, 2009) |
|  | BEDTools | 2.25.0 | Conda | | (Quinlan and Hall, 2010) |
|  | BCFtools | 1.9 | Conda | | (Li, 2011) |
|  | VCFtools | 0.1.16 | Conda | | (Danecek *et al.*, 2011) |
|  | PySimpleGUI* | 4.40.0 | Conda | | (*PySimpleGUI*, no date) |
|  | Snakemake** | ≥ 5.32.1 | Conda | | (Mölder *et al.*, 2021) |

**Supplementary Table 2.** List of tools and corresponding versions that DNAscan2 requires for each pipeline stage. ‘General’ refers to tools which are necessary for the basic execution of DNAscan2. The ‘Installation Method’ column lists the easiest way to obtain the software to avoid software version clashes. *This is only required if you want to run DNAscan2 as a standalone application and not as a command-line tool. **This is only required if you want to deploy DNAscan2 on a high performance computing facility or execute the workflow in a multi-sample parallel fashion.

|  | DNAscan | DNAscan2 |
| --- | --- | --- |
| *SNV/Indel Calling*  - Total called variants  - Total called and filtered variants  - Number of filtered non-synonymous variants  - Number of filtered synonymous variants | 29192123  4285989  10165 (0.237%)  21304 (0.497%) | 30000110  4611498  10727 (0.250%)  22285 (0.483%) |
| *Structural Variant Calling*  - Total called variants   - Deletions - Insertions - Inversions - Duplications - Translocations | 28831  8478  6455  3410  4031  6457 | 56970  16069  6189  7280  9285  18147 |
| *MEI Calling*  - Total called variants   - ALU - SVA - LINE1 | N/A  N/A  N/A  N/A | 1694  1287  148  259 |
| *Expansion/Repeat Scanning*  - Total called and genotyped expansions  - Total Identified STR loci | 37  N/A | 37  3043 |

**Supplementary Table 3.** Comparison of the variant calling power of DNAscan and DNAscan2 for several variant classes. ‘N/A’ represents tasks that could not be performed in DNAscan due to lack of available software. Synonymous and non-synonymous SNV and indel variants were obtained from the corresponding refGene annotation classification.

|  | Pipeline Stage | Pipeline Step | Elapsed Time  (hh:mm:ss) | CPU Time  (hh:mm:ss) | RAM Usage (Gb) | MaxDiskRead | MaxDiskWrite |
| --- | --- | --- | --- | --- | --- | --- | --- |
| DNAscan | Analysis | SNV/Indel Calling | 07:48:52 | 32:10:31 | 34.34 | 144046 | 71073 |
|  |  | SV Calling | 02:46:41 | 11:06:44 | 0.69 | 827571 | 338 |
|  |  | MEI Calling | - | - | - | - | - |
|  |  | Known Expansion | 00:19:48 | 01:19:10 | 9.55 | 23669 | 1 |
|  |  | STR | - | - | - | - | - |
|  | Annotation | SNV/Indel | 03:06:26 | 12:25:44 | 7.74 | 402697 | 71254 |
|  |  | SV + MEI | - | - | - | - | - |
|  | Report Generation | SNV/Indel | 01:24:58 | 05:39:51 | 0.02 | 1953279 | 8 |
|  |  | SV + MEI | - | - | - | - | - |
| DNAscan2 | Analysis | SNV/Indel Calling | 03:54:10 | 15:36:41 | 1.00 | 81794 | 9294 |
|  |  | SV Calling | 33:32:53 | 133:53:30 | 5.70 | 1222044 | 55738 |
|  |  | MEI Calling | 05:53:05 | 23:32:20 | 8.75 | 2710405 | 38612 |
|  |  | Known Expansion | 00:19:48 | 01:19:10 | 9.55 | 23669 | 1 |
|  |  | STR | 00:44:51 | 02:59:24 | 0.66 | 28505 | 30 |
|  | Annotation | SNV/Indel | 03:00:48 | 12:03:12 | 6.78 | 353781 | 35076 |
|  |  | SV + MEI | 00:47:15 | 03:08:58 | 13.54 | 8653 | 8386 |
|  | Report Generation | SNV/Indel | 01:22:17 | 05:29:06 | 0.02 | 1840523 | 10 |
|  |  | SV + MEI | 00:11:52 | 00:47:27 | 15.00 | 980 | 2331 |

**Supplementary Table 4.** Comparison of the variant calling power of DNAscan and DNAscan2 for several variant classes. ‘N/A’ represents tasks that could not be performed in DNAscan due to lack of available software. Synonymous and non-synonymous SNV and indel variants were obtained from the corresponding refGene annotation classification.

| Gene Name | Chromosome | Sources of Identification | | | Reference |
| --- | --- | --- | --- | --- | --- |
|  |  | ***Manual Review*** | ***ALSoD*** | ***ClinVar*** |  |
| DNAJC7 | 17q21.2 | 🗸 |  |  | (Farhan *et al.*, 2019) |
| ERBB4 | 2q34 | 🗸 |  |  | (Takahashi *et al.*, 2013) |
| ANXA11 | 10q22.3 |  | 🗸 |  | (Smith *et al.*, 2017, p. 11) |
| ARPP21 | 3p22.3 |  | 🗸 |  | (Cooper-Knock *et al.*, 2019) |
| ATXN1 | 6p22.3 |  | 🗸 |  | (Tazelaar *et al.*, 2020) |
| C21orf2 | 21q22.3 |  | 🗸 | 🗸 | (van Rheenen *et al.*, 2016) |
| CAV1 | 7q31.2 |  | 🗸 |  | (Cooper-Knock *et al.*, 2020) |
| CAV2 | 7q31.2 |  | 🗸 |  | (Cooper-Knock *et al.*, 2020) |
| CCNF | 16p13.3 |  | 🗸 | 🗸 | (Williams *et al.*, 2016) |
| DNMT3A | 2p23.3 |  | 🗸 |  | (Chestnut *et al.*, 2011) |
| DNMT3B | 20q11.21 |  | 🗸 |  | (Chestnut *et al.*, 2011) |
| ENAH | 1q42.12 |  | 🗸 |  | (Cirulli *et al.*, 2015) |
| EPHA3 | 3p11.1 |  | 🗸 |  | (Uyan *et al.*, 2013) |
| ERLIN1 | 10q24.31 |  | 🗸 | 🗸 | (Tunca *et al.*, 2018) |
| GLT8D1 | 3p21.1 |  | 🗸 |  | (Cooper-Knock *et al.*, 2019) |
| GPX3 | 5q33.1 |  | 🗸 |  | (Benyamin *et al.*, 2017) |
| KIF5A | 12q13.3 |  | 🗸 |  | (Nicolas *et al.*, 2018) |
| MOBP | 3p22.1 |  | 🗸 | 🗸 | (van Rheenen *et al.*, 2016) |
| NEFL | 8p21.2 |  | 🗸 | 🗸 | (Benatar *et al.*, 2018) |
| NEK1 | 4q33 |  | 🗸 |  | (Kenna *et al.*, 2016) |
| PGRN | 17q21.31 |  | 🗸 |  | (Philips *et al.*, 2010) |
| PNPLA6 | 19p13.2 |  | 🗸 | 🗸 | (Pensato *et al.*, 2020) |
| RFTN1 | 3p24.3 |  | 🗸 |  | (Zhai *et al.*, 2009) |
| SCFD1 | 14q12 |  | 🗸 | 🗸 | (van Rheenen *et al.*, 2016) |
| TIA1 | 2p13.3 |  | 🗸 | 🗸 | (Mackenzie *et al.*, 2017) |
| UBQLN1 | 9q21.32 |  | 🗸 |  | (Wang, Tatman and Monteiro, 2020, p. 1) |
| VRK1 | 14q32.2 |  | 🗸 | 🗸 | (Nguyen *et al.*, 2015) |
| CAPN14 | 2p23.1 |  |  | 🗸 | (Dekker *et al.*, 2019) |
| CHRNA3 | 15q25.1 |  |  | 🗸 | (Sabatelli *et al.*, 2009) |
| CHRNB4 | 15q25.1 |  |  | 🗸 | (Sabatelli *et al.*, 2009) |
| CNTF | 11q12.1 |  |  | 🗸 | (*Serum level of CNTF is elevated in patients with amyotrophic lateral sclerosis and correlates with site of disease onset - Laaksovirta - 2008 - European Journal of Neurology - Wiley Online Library*, no date) |
| CYLD | 16q12.1 |  |  | 🗸 | (Dobson-Stone *et al.*, 2020) |
| DDX20 | 1p13.2 |  |  | 🗸 | (Cacciottolo *et al.*, 2019) |
| DYNC1H1 | 14q32.31 |  |  | 🗸 | (Scarlino *et al.*, 2020) |
| ELP3 | 8p21.1 |  |  | 🗸 | (Bento-Abreu *et al.*, 2018) |
| EWSR1 | 22q12.2 |  |  | 🗸 | (Couthouis *et al.*, 2012) |
| GARS1 | 7p14.3 |  |  | 🗸 | (Corcia *et al.*, 2019) |
| GLE1 | 9q34.11 |  |  | 🗸 | (Kaneb *et al.*, 2015) |
| MFN2 | 1p36.22 |  |  | 🗸 | (Wang *et al.*, 2018, p. 2) |
| MPZ | 1q23.3 |  |  | 🗸 | (Bisogni *et al.*, 2021) |
| NIPA1 | 15q11.2 |  |  | 🗸 | (Blauw *et al.*, 2012) |
| PLEKHG5 | 1p36.31 |  |  | 🗸 | (Gonzalez-Quereda *et al.*, 2021) |
| PON1 | 7q21.3 |  |  | 🗸 | (Verde *et al.*, 2019) |
| PON3 | 7q21.3 |  |  | 🗸 | (Saeed *et al.*, 2006) |
| SCN11A | 3p22.2 |  |  | 🗸 | (Castoro *et al.*, 2018) |
| SLC52A3 | 20p13 |  |  | 🗸 | (Johnson *et al.*, 2012) |
| SYNE1 | 6q25.2 |  |  | 🗸 | (Naruse *et al.*, 2020) |
| TRPV4 | 12q24.11 |  |  | 🗸 | (Pensato *et al.*, 2020) |
| UNC13A | 19p13.11 |  |  | 🗸 | (Diekstra *et al.*, 2012) |

**Supplementary Table 5.** New additions and/or reclassifications in the ALSgeneScanner database. Breakdown of identified gene sources are as follows: 2 manual review, 16 ALSoD, 9 ALSoD and ClinVar, and 22 ClinVar.
